# Vascular bundle sheath and mesophyll regulation of leaf water balance in response to chitin

**DOI:** 10.1101/337709

**Authors:** Ziv Attia, Ahan Dalal, Menachem Moshelion

## Abstract

Plants can detect pathogen invasion by sensing pathogen-associated molecular patterns (PAMPs). This sensing process leads to the induction of defense responses. Most PAMP mechanisms of action have been described in the guard cells. Here, we describe the effects of chitin, a PAMP found in fungal cell walls, on the cellular osmotic water permeability (*P*_f_) of the leaf vascular bundle-sheath (BS) and mesophyll cells and its subsequent effect on leaf hydraulic conductance (*K*_leaf_).

The BS is a parenchymatic tissue that tightly encases the vascular system. BS cells have been shown to control *K*_leaf_ through changes in their *P*_f_, for example, in response to ABA. It was recently reported that, in Arabidopsis, the chitin receptors chitin elicitor receptor kinase 1 (CERK1) and LYSINE MOTIF RECEPTOR KINASE 5 (LYK5) are highly expressed in the BS, as well as the neighboring mesophyll. Therefore, we studied the possible impact of chitin on these cells.

Our results revealed that both BS cells and mesophyll cells exhibit a sharp decrease in *P*_f_ in response to chitin treatment. In addition, xylem-fed chitin decreased *K*_leaf_ and led to stomatal closure. However, an *Atlyk5* mutant showed none of these responses. Complimenting *AtLYK5* specifically in the BS cells (using the SCARECROW promoter) and transient expresion in mesophyll cells each resulted in a response to chitin that was similar to that observed in the wild type. These results suggest that BS and mesophyll cells each play a role in the perception of apoplastic chitin and in initiating chitin-triggered immunity.

**Significance Statement:** PAMP perception by plant receptors triggers various defense responses important for plant immunity. Here we provide new insights into a topic that has received a great deal of previous attention, revealing that a chitin immune response is present in additional leaf tissues other than the stomata. Chitin perception by the bundle sheath cells enwrapping the whole leaf vascular system decrease its cellular osmotic permeability and leaf hydraulic conductance. This in turn, leads to hydraulic signals being sent to the stomata and regulates whole-leaf water balance in response to chitin application and, perhaps, during fungal infection. Emphasizing the dynamic role of the BS in chitin-sensing and water balance regulation.

## INTRODUCTION

Plants are constantly exposed to various microorganisms. Pathogenic microorganism challenge may lead to a compatible interaction (successful infection leading to disease) or an incompatible interaction (successful plant defense). However, generally, there is a continuum of susceptibility/resistance to any given pathogen ^1^. Plants have evolved immune systems to defend against microbial infections ^2^. Immunity is initiated by the perception of pathogen-associated molecular patterns (PAMPs), which include fungal chitin, bacterial flagellin, the elongation factor Tu (EF-Tu), peptidoglycan and other substances. The detection and sensing of PAMPs is mediated by pattern recognition receptors in the plant’s plasma membranes ^3,4^ and the stimulation of those receptors leads to PAMP-triggered immunity (PTI) ^5^. In the case of pathogenic fungi, the fungal cell wall plays an important role in the plant–fungus interaction. The cell wall is the first part of the pathogen to make physical contact with plant cells, which can recognize several fungal cell-wall components as PAMPs. This recognition activates plant immune responses ^6,7^. Chitin is a major component of fungal cell walls ^8^ and is not found in plants. It acts as a general elicitor for the plant immune response ^9–11^. Fragments of chitin, Λ/-acetylchitooligosaccharides, have been shown to act as potent PAMP signals in various plant species, including *Arabidopsis thaliana* ^12–16^. Challenged plant cells secrete hydrolytic enzymes, such as chitinases, to target the fungal cell-wall constituents and affect the integrity of the fungal cell wall ^17,18^. This activity serves a dual function for the host; the hydrolytic activity may lead to fungal cell collapse resulting in the arrest of pathogen ingress and may also release PAMP molecules that stimulate further PTI responses ^19^.

Chitin-triggered immunity is generally characterized by the induction of various defense responses such as rapid and transient membrane depolarization ^20^, the generation of reactive oxygen species (ROS) ^21^ and the closing of stomata ^22–25^.

Lysine motif (LysM)-containing proteins were previously shown to be involved in plants’ recognition of chitin. Arabidopsis has five members of the lysine-motif receptor-like kinase *(LYKs)* family *(AtCERKULysM RLK1/AtLYK1* and *AtLYK2-5)* ^26^. Consistent with reports that *cerk1-mutant* plants exhibit heightened susceptibility to fungal pathogens ^27^ and the fact that *AtCERK1* can be precipitated by binding to chitin beads ^12,28,29^, *AtCERK1* has been reported to be the primary chitin receptor and to be essential for chitin-induced signaling ^12,28^. Although the X-ray crystal structure of the ectodomain of *AtCERK1* indicates that it is a chitin-binding protein, a calorimetric analysis found that it had a low binding affinity for that substance ^30^. Recently Cao et al. (2014) confirmed that *AtCERK1, AtLYK4* and *AtLYK5,* but not *AtLYK2* or *AtLYK3* are involved in chitin-induced signaling. However, *AtLYK5* binds to chitin with a much higher affinity than *AtCERK1.* Those researchers also suggested that *AtLYK5* is the primary receptor for chitin. Moreover, *AtLYK5* expression was found to be higher than that of *AtCERK1* and *AtLYK4* in the roots and AtLYK5 was found to be a membrane protein ^31^. This might suggest that *AtLYK5* plays a role in defense signaling in the presence of soil-borne vascular pathogens. In addition, it was suggested that *LYM2* mediates a decrease in cell-to-cell connectivity via plasmodesmata in the presence of chitin. This reduction in the symplastic pathway via plasmodesmata is a less characterized PAMP-triggered response that occurs independently of the known intracellular signaling pathways used in PTI and might be important for disease resistance ^32^.

While most aerial plant tissues are protected by a layer of cuticle, the stomata provide a direct path of entry to the plant tissues and are particularly vulnerable to pathogen infection ^33^. The restriction of pathogen entry by stomatal closure is one of the PTI responses that can be detected within minutes and is critical for plants’ innate immunity ^22,23^. Nevertheless, pathogenic fungi can be found in the root vasculature, indicating that they have evolved ways to penetrate the endodermis ^34,35^ and target the xylem vascular system. Moreover, diseases caused by soil-borne fungal pathogens are a major constraint for crop yield and yield quality, particularly in intensive cropping systems ^36^. This points to the importance of understanding plant–pathogen interactions in vascular tissues.

Bundle-sheath (BS) cells, which tightly encase the entire vascular system, have been reported to act as the selective barrier between the dead xylem and living mesophyll cells. BS cells were reported to sense abiotic stress signals (e.g., ABA) through the xylem sap and respond to those signals by changing the osmotic water permeability (*P*_f_) of their membranes. This, in turn, regulates leaf radial hydraulic conductance (*K*_leaf_), thereby regulating the movement of water into the leaf mesophyll cells ^37^ In addition, a recent microarray analysis showed that, in Arabidopsis, 45% of the genes that are differentially expressed between BS and mesophyll cells are membrane-related and 20% are transport-related ^38^. This strengthens the idea that BS cells are a point of control for the radial transport of solutes. The transcriptomic data also indicate that *AtLYK5* is expressed at detectable levels in both BS and mesophyll cells, as seen in the normalized log_2_ data (intensity value) ^38^. To act as a checkpoint for the movement of any substance into the mesophyll cells from the vascular tissues, the BS cells must possess intricate mechanisms for sensing any biotic factors that might appear in the vascular tissue, including PAMPS. In this study, we investigated the role of BS cells in chitin-sensing and the regulation of *K*_leaf_ through the modulation of the permeability of the BS cell membrane to water.

## RESULTS

### Chitin Inhibited the *P*_f_ of BS Cells and Mesophyll Cells of Arabidopsis

The transcriptomic data sets of Wigoda et al. (2017) revealed high levels of expression of both *AtCERK1* and *AtLYK5* [10th (top) decile of expression]. Those genes play important roles in chitin sensing and signaling machinery, in both mesophyll cells and BS cells, which motivated us to examine the osmotic water permeability (*P*_f_) of both of those type of cells in response to chitin. GFP-labeling of BS cells was used to distinguish those cells from the (unlabeled) mesophyll cells. [Protoplasts were prepared from SCR::GFP plants and both the cell types were exposed to chitin (see Materials and Methods).]

We first confirmed the sensitivity to chitin of the SCR::GFP plants by evaluating their stomatal closure upon treatment with 0.1 mg/mL chitin (**Supplementary Fig. S2**). Subsequently, we extracted protoplasts from those plants and measured their Pf. During the hypotonic wash (with or without chitin), the volumes of the protoplasts of mesophyll cells and BS cells both increased (**Fig. 1a, b**) in response to the change in the osmotic concentration (C_out_) of the bath (**Fig. 1c, d**). Chitin application resulted in significant reductions in the initial *P*_f_ for both mesophyll cells (37%) and BS cells (49%), relative to their respective untreated controls (**Fig. 1e, f**). However, compared to the mesophyll cells, there was a significant decrease in the *P*_f_ slope for the BS cells (70%) upon chitin treatment (**Fig. 1g, h**). In comparison, there was only a 25% decrease in the *P*_f_ slope of the mesophyll cells. (*P*_f_ slope refers to the change in cell volume-change rate during a change in membrane permeability.) Based on the protoplast assay, at the cellular level, BS cells are significantly more sensitive to chitin than mesophyll cells are, as illustrated by the approximately linear decrease in their *P*_f_ as their volume increased. To ensure that the observed permeability change was not due to any physical and/or permanent damage to the membrane during protoplast isolation or chitin treatment, we measured the radii of the cells before the hypotonic wash and after recovery from that wash. We did not find any significant differences in cell radius between the treated and untreated conditions for BS cells or mesophyll cells. That finding indicates that the cells maintained their ability to act as perfect osmometers (**Supplementary Fig. S3**).

**Figure 1:**
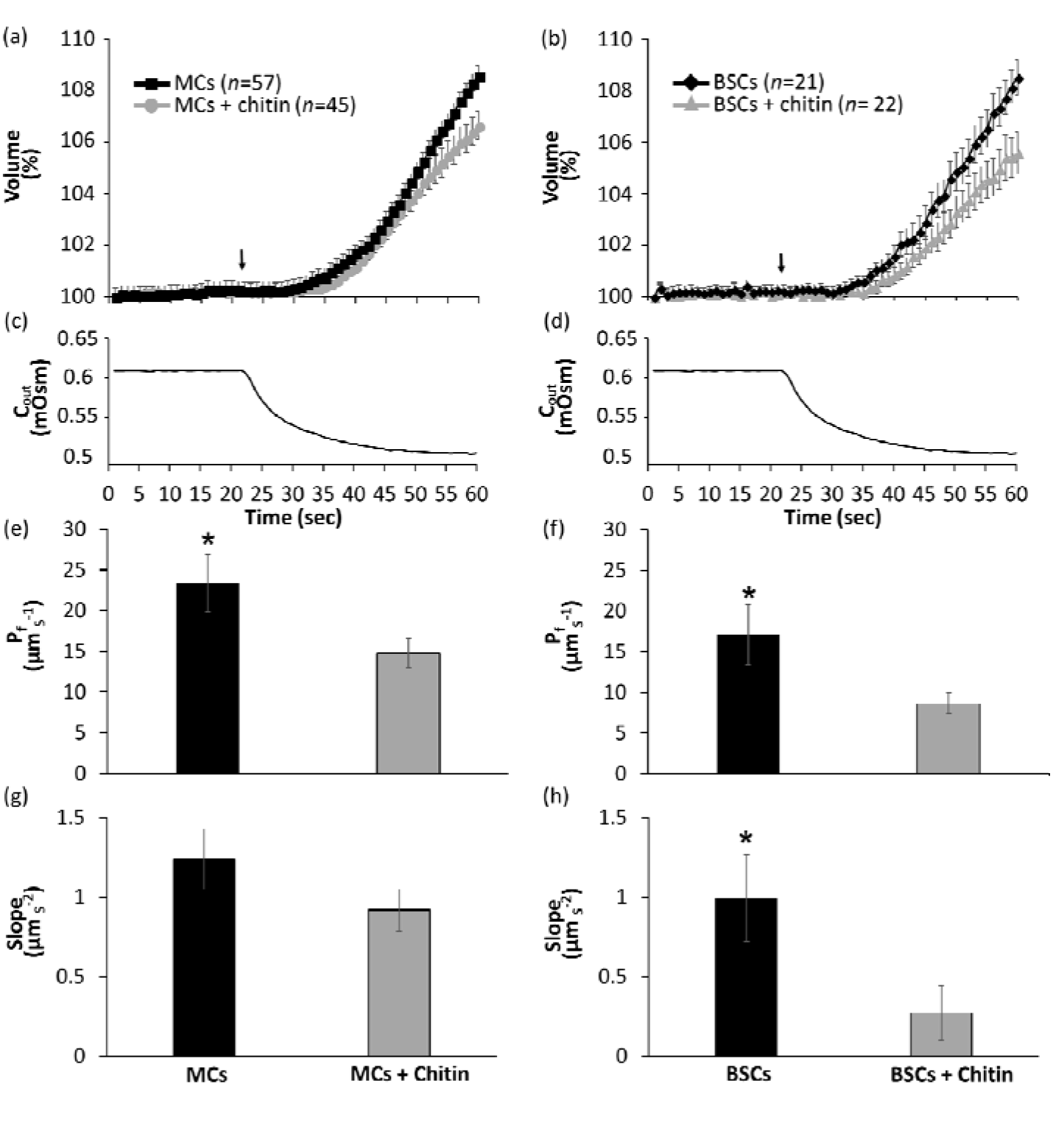
The effect of chitin treatment on the membrane osmotic water permeability *(P_f_)* of MCs and BSCs from WT Arabidopsis. Time course (60 sec) of the volume increase of (a) Mesophyll and (b) bundle sheath protoplasts upon exposure to hypotonic solution with or without chitin, the arrow indicates onset of bath flush. (c,d) Time course of the osmotic concentration change in the bath (C_out_) during the hypotonic wash for mesophyll and bundle sheath protoplasts. (e,f) Mean *P*_f_ of mesophyll and bundle sheath protoplasts during chitin application. (g,h) Mean slope of mesophyll and bundle sheath protoplasts during chitin application. Data is shown as means ±SE. Asterisks represent significant differences between treatments (Student’s t-test, P < 0.05).

In order to further understand the effect of chitin on water balance at the cellular level, we exposed the isolated mesophyll protoplasts from *lyk5* mutant plants to chitin and measured their *P*_f_. Upon exposure to the hypotonic wash, no significant differences in cell volume, *P*_f_ or *P*_f_ slope were observed between the treated and untreated cells (**Fig. 2a, c, e, g**). When we complemented lyk5-mutant mesophyll cells with *AtLYK5+GFP,* the hypotonic wash caused an increase in protoplast volume among both treated and untreated protoplasts (**Fig. 2b, d**). However, chitin application resulted in a significant (73%) reduction in the *P*_f_ of the complemented mesophyll cells (**Fig. 2f**) and a decrease in their *P*_f_ slope (86%), relative to the untreated controls (**Fig. 2h**). Again, to confirm that the permeability change was not due to any physical and/or permanent damage done to the membrane during protoplast isolation, PEG transformation or chitin treatment, we measured the radii of the cells before hypotonic challenge and after recovery from the hypotonic challenge. We did not find any significant differences between the cell radii of the treated and untreated, transformed and untransformed cells (**Supplementary Fig. S4**).

**Figure 2.**
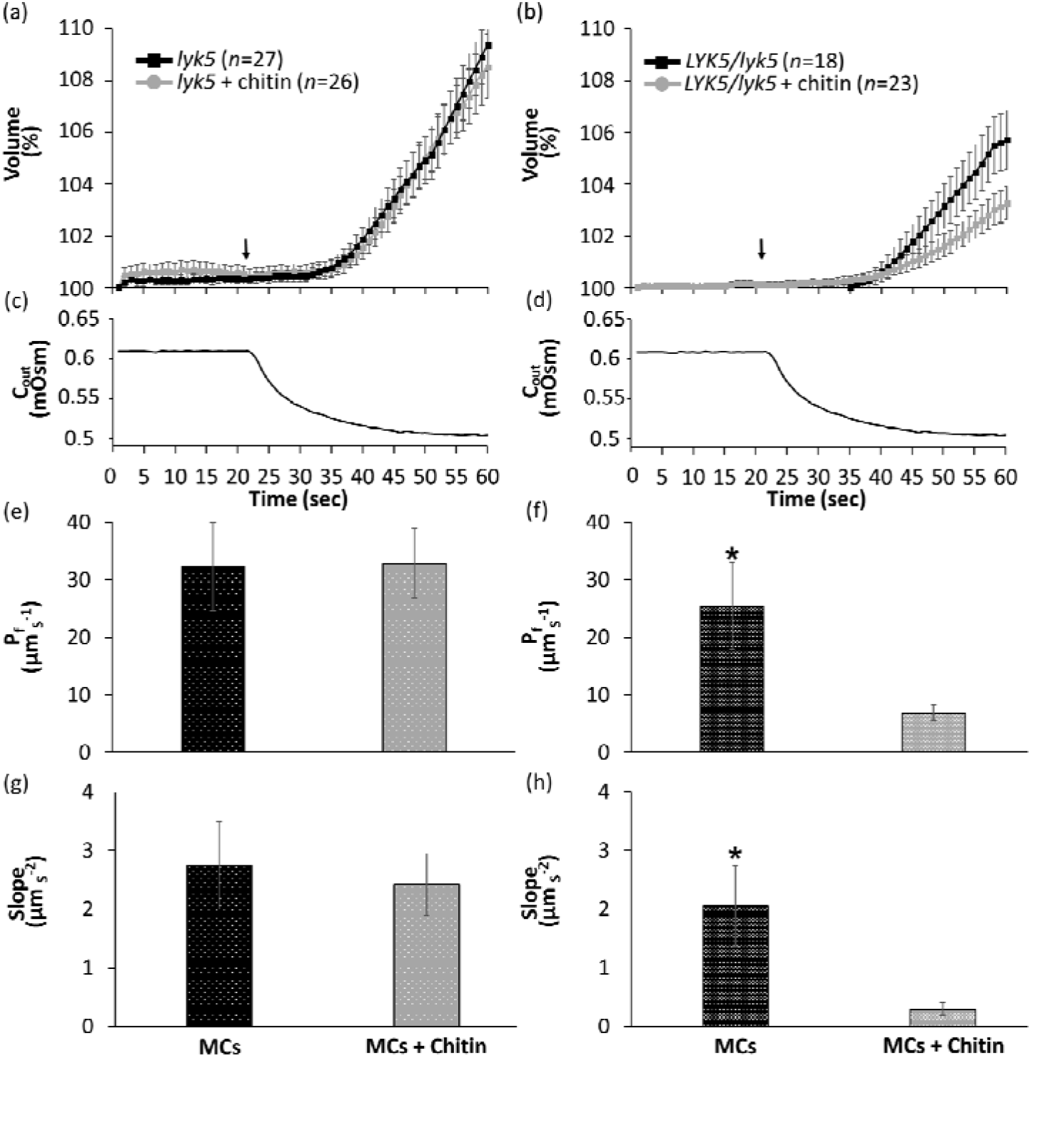
The effect of chitin treatment on the membrane osmotic water permeability (*P*_f_) of mesophyll cells from *lyk5* mutants and *AtLYK5*-complemented *lyk5* mutants. Time course (60 s) of the volume increase of (a) untransformed mesophyll and (b) transformed mesophyll protoplasts upon exposure to hypotonic solution with or without chitin. Arrows indicate the start of the bath treatment. (c, d) Time course of the osmotic concentration change in the bath (C_out_) during the hypotonic wash for untransformed and transformed mesophyll protoplasts. (e, f) Mean *P*_f_ of untransformed and transformed mesophyll protoplasts during chitin application. (g, h) Mean slope of untransformed and transformed mesophyll protoplasts during chitin application. Data are shown as means ± SE. Asterisks represent significant differences between treatments (Student’s t-test, P < 0.05).

### Xylem-Fed Chitin Suppressed the *K*_leaf_ of Arabidopsis

The role of BS cells in the regulation of the radial flow of water through the tight apoplastic barrier between the xylem and the mesophyll cells, the greater sensitivity of BS cells to chitin (**Fig. 1**) and the expression of the chitin receptor *AtLYK5* in BS cells, all suggest that BS cells are involved in the regulation of *K*_leaf_ in response to chitin. To further explore this matter, we evaluated the effect of chitin application on the *K*_leaf_ of the WT, *lyk5* mutants and BS-specific complemented plants (SCR::LYK5/lyk5) in which chitin sensing exists solely in the BS cells. As the BS cells act as a barrier to small solutes ^37^, the detached-leaf approach allowed us to feed the chitin directly to the xylem via the petiole, enabling direct contact between the BS cells and the applied chitin ^38–40^. In order to ensure the reliability of the chitin treatment, we used two different methods of treatment, a submerged-leaf method and a petiole-feeding method, and used qPCR to examine the quantitative expression of the chitin-induced marker genes *AtWRKY29* and *AtWRKY30* ^31^ in the WT. In the case of the submerged-leaf treatment, the expression of both of these genes in the treated plants was significantly higher than the expression levels observed in the untreated control (**Fig. 3a, b**). However, with the petiole-fed treatment, only *AtWRKY30* expression was significantly higher in the treated plants, as compared to the untreated control (**Fig. 3c, d**). This confirms the credibility of our petiole-feeding experimental setup, which allows for chitin perception by the leaf.

**Figure 3.**
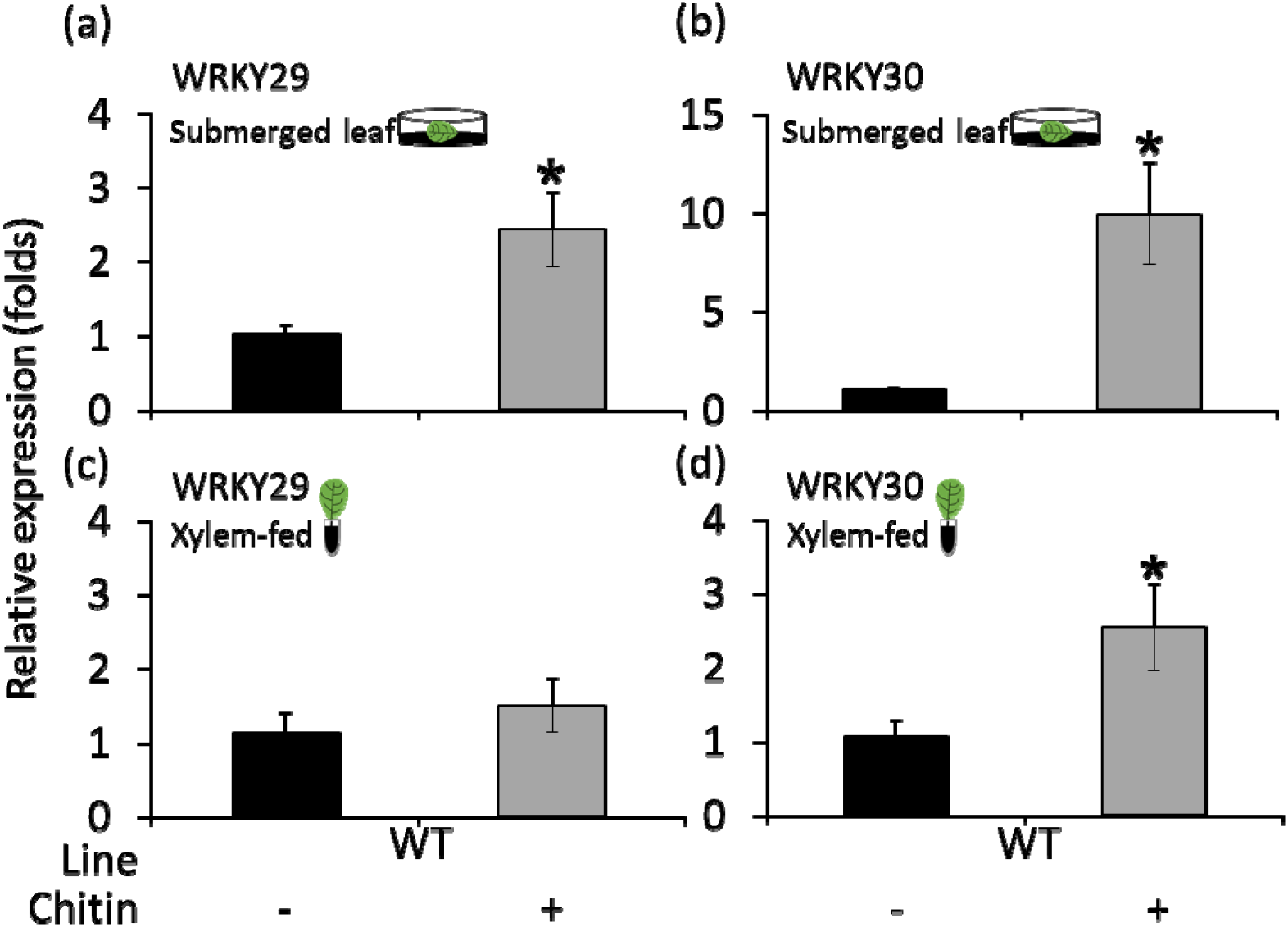
Relative expression of WRKY29 and WRKY30 genes in the whole leaf. *AtWRKY29 (At4g23550)* and *AtWRKY30 (At5g24110)* gene expression was analyzed using qRT-PCR in either (a-b) submerged or (c-d) xylem-fed WT leaves. Data are shown as means ± SE of at least three independent experiments. An asterisk above the column indicates a significant difference between treatments (t-test, *P* < 0.05).

In order to validate chitin insensitivity in tissues other than BS cells in the complemented SCR::LYK5/lyk5 plants, we analyzed chitin-mediated stomatal closure in epidermal peels. Neither SCR::LYK5/lyk5 nor the *lyk5* mutant showed any response to chitin; whereas the WT exhibited a significant decrease in stomatal aperture in response to chitin treatment. As a positive control, we used 1 μM ABA, which resulted in a significant decrease in stomatal aperture in all of the different plants (i.e., WT, the *lyk5* mutant and the SCR::LYK5/lyk5 plants; **Fig. 4**). This phenotype was observed in all three of the independent SCR::LYK5/lyk5 transgenic lines tested (**Supplementary Fig. S5**).

**Figure 4.**
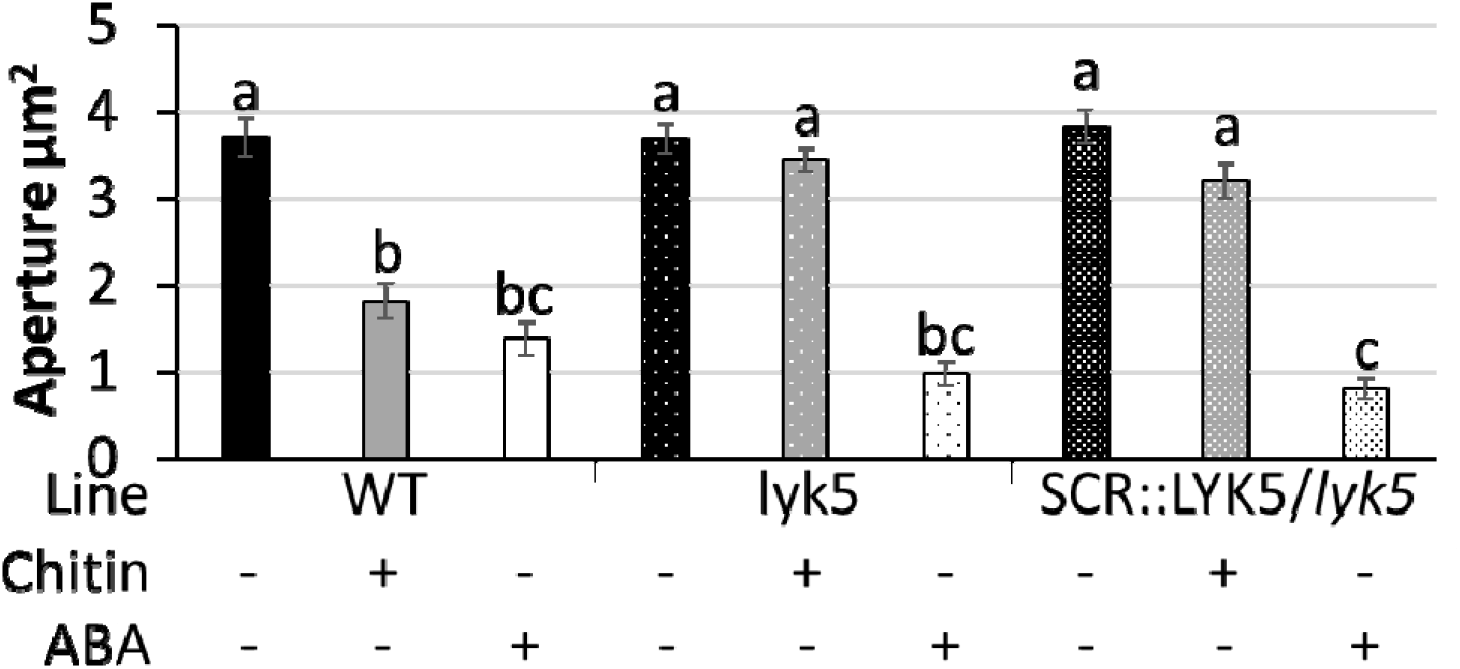
Chitin application did not decrease stomatal aperture in the epidermal peels of the *lyk5* mutant or the SCR::LYK5/lyk5 plants. Images of abaxial epidermis were taken from leaves of 8-week-old Col-0 plants, the *lyk5* mutant and SCR::LYK5-complemented *lyk5* plants. ABA-treated plants were used as a control. Stomatal aperture was measured after an incubation of 1.5 h. Data are shown as means ± SE. Different letters above the columns indicate significant differences between treatments, according to Tukey’s HSD test (*P* < 0.05). (*n* >60)

Petiole-fed chitin induced a 56% decrease in *K*_leaf_ among the WT plants, no change in *K*_leaf_ among the *lyk5* mutants and a 57% decrease in *K*_leaf_ among the BS-specific complemented *lyk5* mutants, as compared to their respective untreated controls (**Fig. 5a**). When compared to their respective untreated controls, the treatment reduced *E* by ~20% among the WT plants, caused no change in *E* among the *lyk5* mutants and caused a 30% decrease in *E* among the BS-specific complemented *lyk5* mutants (**Fig. 5b**). Chitin treatment also decreased Ψ_leaf_ by ~50% among the WT plants, caused no change in the Ψ_leaf_ of *lyk5* mutants and caused a 43% decrease in the Ψ_leaf_ of the BS-specific complemented *lyk5* mutants, as compared to their respective untreated controls (**Fig. 5c**). This phenotype was observed in all three independent SCR::LYK5/lyk5 transgenic lines and among the SCR::GFP plants (**Supplementary Fig. S6, S7**).

**Figure 5.**
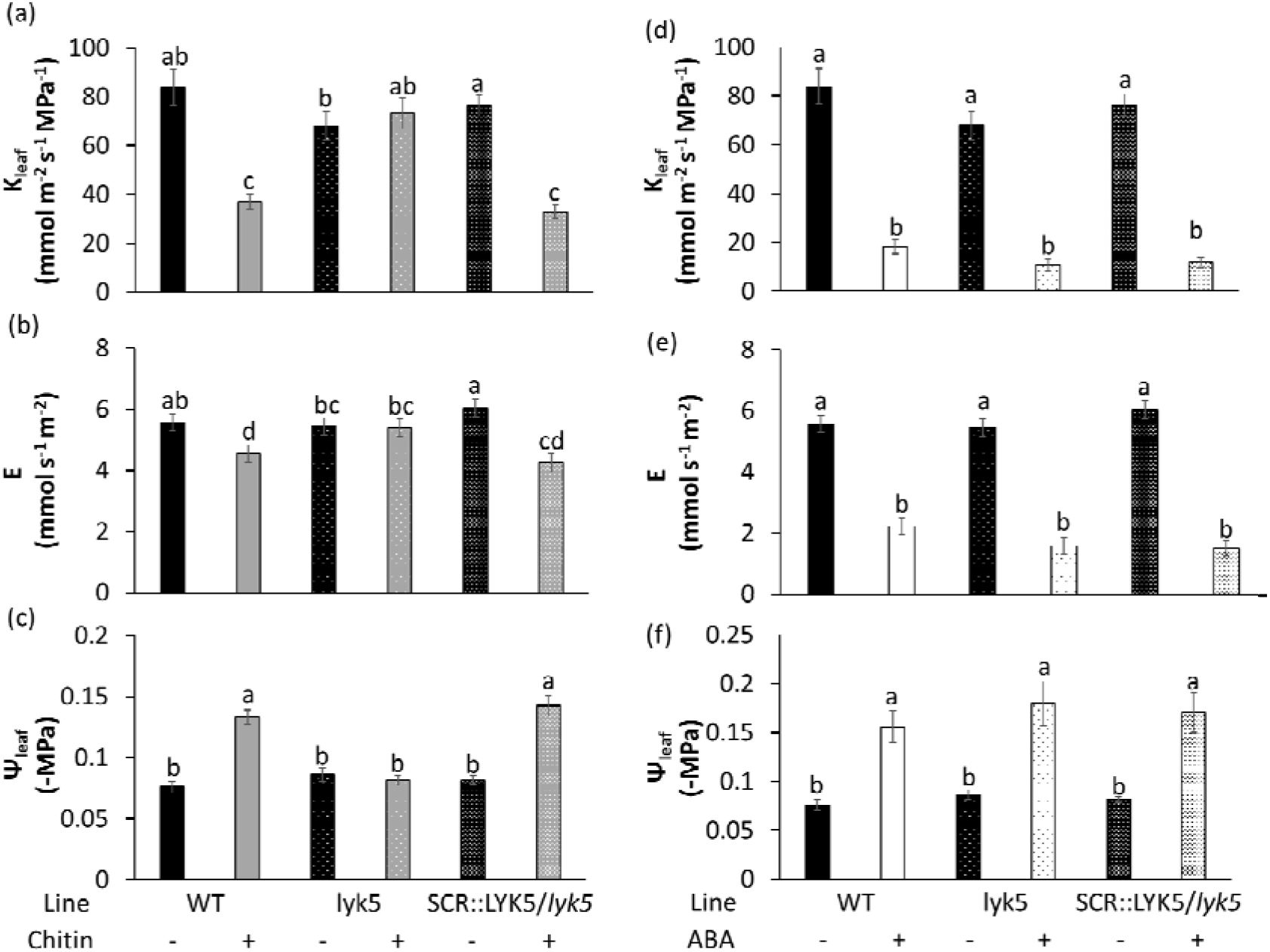
Complementation of *lyk5* in BS cells led to the recovery of leaf hydraulic sensitivity to chitin. (a) Leaf hydraulic conductance, *K*_leaf_, (b) transpiration rate, *E*, and (c) water potential, Ψ_leaf_ of WT, *lyk5* mutant and BS-specific LYK5-complemented plants after 2–4 h of the xylem-fed chitin treatment. One μM ABA was used as a positive control to validate the experimental setup: (d) *K*_leaf_, (e) *E* and (f) Ψ_leaf_. Different letters above the columns represent significant differences between treatments (Tukey’s HSD test, *P* < 0.05). Data are means (±SEs) from at least three independent experiments.

To sum up, the chitin receptor mutant *lyk5* revealed no significant changes in *K*_leaf_, *E* or K_leaf_ in response to chitin, as compared to the untreated controls. However, BS-specific complementation of the mutant resulted in plants that were sensitive to chitin treatment, much like the WT. As a positive control, we used 1 μM ABA, which reduced *K*_leaf_, *E* and ψ_leaf_ in all of the different plants (i.e., WT, *lyk5* mutant and the BS-specific complemented mutants; **Fig. 5 d, e, f**). This reduction in response to petiole-fed ABA, which was observed among all of the different plants, is similar to that observed among WT Arabidopsis in earlier studies ^37^.

## DISCUSSION

Perception of PAMPs and the subsequent PTI are the first line of a multilayered defense system in plants ^41,42^. Previous studies have shown that PTI immune responses occur upon the perception of chitin by *AtCERK1* and *AtLYK5,* initiating the well-documented hallmarks of PTI responses, such as the production of ROS ^13,43^, the induction of defense ^31^, marker genes and stomatal closure ^22–25^. In this work, we studied the role of chitin in internal tissues and revealed its role in regulating *P*_f_, which affects leaf hydraulic conductance and water balance.

Both *AtCERK1* and *AtLYK5* RNA are highly expressed in both mesophyll cells and BS cells ^38^. Accordingly, our data show that both isolated BS cells and mesophyll cells sense chitin independently, leading to a sharp decrease in Pf. While both BS cells and mesophyll cells responded to chitin treatment, the BS cells exhibited a ~50% drop in *P*_f_ and a 70% drop in the *P*_f_ slope, indicating their greater sensitivity to / reactivity with chitin (**Fig. 1**). In addition, complementation of the *Atlyk5* mutant with *AtLYK5* resulted in a significant decrease in *P*_f_ in response to chitin application, as compared with the non-sensing mutant (**Fig. 2**). This indicates that the observed reduction in *P*_f_ occurs upon the autonomous perception of chitin at the cellular level.

Stomatal immunity or the closure of stomata upon PAMP recognition has attracted considerable attention in recent years. Like other known PAMPs, chitin induces stomatal closure. In fact, the restriction of microbial entry by stomatal closure, known as pre-invasive or stomatal immunity, is one of the PTI responses that can be detected within a very short period of time (**Fig. 4**) ^22^. Stomatal-closure responses to ABA and PAMPs have been shown to involve an increase in guard-cell water permeability (higher *P*_f_). Mediated by the activity of aquaporins, this change in guard-cell *P*_f_ has been shown to result in a decrease in guard-cell volume and subsequent stomatal closure ^44,45^. Previous work and our results show that the application of both chitin and ABA leads to a decrease in the *P*_f_ of BS cells ^37,45^ (**Fig. 1**). Thus, BS cells and stomata seem to exhibit antagonistic *P*_f_ responses to PAMPs and ABA (biotic and abiotic stress signals) ^37,45^.

BS cells have been shown to play a role in creating a xylem–mesophyll hydraulic barrier under water-stress conditions ^37,46^, regulating the apoplastic xylem flow into the leaf via the BS cell transmembrane pathway and directly affecting K_leaf_ by regulating their *P*_f_ and aquaporin activity ^47^ Indeed, xylem-feeding with chitin resulted in significant decreases in *K*_leaf_, *E* and Ψ_leaf_. Our results suggest that the reduction in Ψ_leaf_ observed following the application of xylem-fed chitin (**Fig. 5c**) resulted from a change in the balance between water influx and efflux [i.e., inhibition of the movement of water into the leaf mesophyll via a radial trans-BS (xylem-to-mesophyll) pathway]. This, in turn, reduced the *K*_leaf_ mainly by decreasing water vapor efflux via stomatal closure (i.e., by decreasing E; **Fig. 5b**) in a manner similar to ABA ^37^ The complementation of the *Atlyk5* mutation by SCR::LYK5 supports the existence of this vein-to-stomata hydraulic signal, as the expression of *AtLYK5* solely in the BS of the mutant led to the complete recovery of the *K*_leaf_ of the *lyk5* mutant (**Fig. 5a-c**), despite the fact that in epidermal peels of the SCR::LYK5 the stomata did not response to chitin treatment (**Fig. 4**). The fact that BS cell chitin-sensing and hydraulics control the whole-leaf water balance, supports the existence of a tight hydraulic connection between the BS and the mesophyll ^37,39,47,48^, as well as feed-forward regulation of the mesophyll and stomata. Nevertheless, the reductions observed in these physiological parameters could also be attributed, at least partially, to the mesophyll cells, as the BS can indirectly control mesophyll hydraulic conductance ^47,49^.

The results of this study lead us to ask: What are the roles and advantages of BS and mesophyll chitin-sensing for the plant? While many xylem-invading vascular wilt pathogens are soil-borne and enter their hosts through wounds in the roots or cracks that appear at the sites of lateral root formation, some vascular-wilt pathogens enter plants through the stomata and hydathodes ^50^. While stomatal closure restricts pathogen entry ^22,23^, the hydathodes are non-regulated openings on the leaf margins, which open to a group of thin-walled small cells (epithem) that connects directly with the xylem vessels. Moreover, guttation fluid is enriched with minerals and contains amino acids and sugars, which can support the development of microorganisms ^51,52^. Regardless of the mechanism that vascular-wilt pathogens use to enter their hosts, these pathogens subsequently colonize the xylem vessels, proliferate and spread ^50^. Previous studies have also shown that the xylem sap of both non-infected Arabidopsis and barley *(Hordeum vulgare)* plants contains chitinases ^52,53^.

In light of all of this, we would like to suggest a speculative hypothesis relating to hydathodes’ sensing of pathogens. When a fungal pathogen invades the xylem, the PAMP molecule chitin is generated in the xylem as a result of chitinase activity ^7^ This is proof of the fact that a line of defense exists in the xylem as an innate immune system or at least during/after a xylem-invading pathogen attack. Once a fungal pathogen manages to penetrate the xylem vessels, the next radial point of entry to the leaf is the BS, which acts as a selective barrier between the dead xylem and the living mesophyll cells. Our finding that BS cells are more sensitive to chitin than mesophyll cells are (**Fig. 1**) suggests that the position of BS cells as the first line of defense directly facing the xylem may expose those cells to pathogens in the xylem stream. Therefore, in response to chitin in the xylem stream, BS cells not only restrict the movement of water into the xylem, to minimize the further entry of pathogens through xylem, but also protect against additional pathogen entry by closing the stomata via the hydraulic signal initiated by the reduction in Ψ_leaf_ (**Fig. 5**) ^37,48^. Moreover, the fact that mesophyll cells express chitinase in their apoplastic cell wall ^52,53^ and the fact that the *P*_f_ of mesophyll cells decreases in response to chitin, suggest that these cells may exhibit immune responses similar to those exhibited by the stomata and BS cells. These immune response may be beneficial in cases of inoculation through wounds or as hypha spread through the leaf. However, that hypothesis needs to be explored in future work.

## Conclusion

Our results provide new insights into a topic that has received a great deal of previous attention, revealing that a PAMP immune response is present in tissues other than the stomata and emphasizing the dynamic role of the BS in chitin-sensing, through its control of *K*_leaf_. These results underscore the importance of the BS’s role in chitin-sensing, which, in turn, leads to hydraulic signals being sent to the stomata and regulates whole-leaf water balance in response to chitin application and, perhaps, during fungal infection.

## MATERIALS AND METHODS

### Plant Materials and Growing Conditions

*Arabidopsis thaliana* Col-0 (WT) and the T-DNA insertion mutant line *lyk5* (SALK_131911C) were used in this study. The mutant line was well characterized for the loss of function of the *LYK5* gene ^31^. We obtained the mutant line from NASC: The European Arabidopsis Stock Centre (http://arabidopsis.info) and used qPCR to screen for homozygosity and test for the loss of the gene (**Supplementary Fig. S1**). Complementation lines in which *LYK5* was expressed in BS cells were obtained by transforming the pML-BART-SCR:: *LYK5* plasmid into the *lyk5* background. Transgenic lines expressing GFP in BS cells were obtained by transforming the pML-BART-*SCR::mGFP5-ER* plasmid into the WT (Col-0) background. The generation of transgenic lines is described below.

All plants were either germinated on MS agar medium [Murashige and Skoog medium including Nitsch vitamins with plant agar (Duchefa Biochemie)] with the seedlings then transferred into potting mixture (Klasmann-Deilmann substrate select special mixture) containing slow-release fertilizer (4g/L; Everris Osmocote Pro 3-4M) or germinated directly in the potting mixture. All plants were grown at 16–25°C (night–day) under short-day conditions (8 h light/16 h dark) with 60–70% humidity and light intensity of 100 to 15 0 μmol m^-2^ s^-1^.

### Plasmid Construction

To generate a plant expression cassette with a constitutive promoter, the pART7 vector was used. That vector consists of cauliflower mosaic virus Cabb B-JI isolate, the 35S promoter and the transcriptional termination region of the *octopine synthase* (*OCS*) gene. For BS-specific expression in plants, we chose to use the SCARECROW (SCR) promoter. The SCR promoter [2360 bp, upstream of the *SCR* gene (TAIR: AT3G54220)] was amplified from the gDNA of Arabidopsis leaves. [gDNA was isolated using the method described by Doyle and Doyle (1987)]. The DNA fragment was cloned into the *XhoI* and *KpnI* sites of pART7 vector to generate a SCR-pART7 vector. A *NotI* site was introduced at the 3’ end of the *XhoI* site in order to eliminate the built-in 35S when the expression cassette under the regulation of SCR promoter was subsequently introduced into the *NotI* site of the binary vector pML-BART for plant transformation.

The *LYK5* (TAIR: AT2G33580) CDS (1995 bp) was amplified from the cDNA of WT plants (cDNA synthesis is described in the RNA Isolation and Gene-Expression Analysis section below) and cloned into the *KpnI* and *XbaI* sites of both pART7 and the SCR-pART7, vector, generating the expression cassettes with 35S promoter and SCR promoter, respectively, and the *OCS* terminator. For constitutive GFP expression in Arabidopsis protoplasts, pSAT1-EGFP-C1 was used ^37,54^ For BS-specific expression in plants, *mgfp5-ER* (GenBank U87974.1) complete CDS was amplified and cloned into the *KpnI* and *XbaI* sites of SCR-pART7 vector, to generate an expression cassette with the *SCR* promoter and the *OCS* terminator. The expression cassettes from pART7 and SCR-pART7 were subsequently introduced into the *NotI* site of pML-BART for plant transformation. All amplifications for cloning were done using Phusion High-Fidelity DNA Polymerases (Thermo Fisher Scientific). The cloned plasmids were sequenced (Hy Laboratories Ltd.) to confirm the accuracy of the constructs. The amplification primers used for cloning are listed in Supplementary Table 1.

### Generation of Transgenic Lines

The recombinant plasmids pML-BART-SCR:*:mGFP5-ER* and pML-BART-SCR::*LYK5* carrying the expression cassette was transfected into *Agrobacterium tumefaciens* GV3101 cells, which were then used to transform 6-week-old WT and *lyk5* Arabidopsis plants using the floral-dip method ^55^. T_0_ seeds were screened for glufosinate resistance (TOKU-E) on MS agar plates. The identity of the *SCR::LYK5* T_1_ plants was confirmed by gDNA PCR and transgene expression was investigated by qPCR (Supplementary Fig. S1), as described in the RNA Isolation and Gene-Expression Analysis section below. The identity of the SCR::GFP T_1_ plants was verified by epifluorescence under an inverted microscope (Nikon eEclipse ts100). All experiments were conducted with the T_2_ generation of plants.

### Chitin Preparation and Treatment

For chitin (from shrimp shells, Sigma-Aldrich) treatment, the stock solution was prepared according to the protocol described by ^56^. Briefly, for 10 mg/mL stock solution, chitin suspended in double-distilled water was autoclaved for 30 min. The solution was then centrifuged and the supernatant was used as the stock solution. Solution with a concentration of 0.1 mg/mL of chitin was used for all of the experiments. The chitin-sensing of the different genotypes was evaluated in terms of stomatal closure and gas-exchange measurements.

### RNA Isolation and Gene-Expression Analysis

Total RNA was isolated from *A. thaliana* leaves using the Plant Total RNA Mini Kit (Geneaid) and treated with DNase I (Geneaid) to avoid genomic DNA contamination. Five μg of total RNA were used for first-strand cDNA synthesis, which was carried out using the qPCRBIO cDNA Synthesis Kit (PCRBIOSYSTEMS).

For transgene-expression analysis, untreated plants were used. To analyze the expression of the chitin-induced marker genes *AtWRKY29* and *AtWRKY30* in the WT, chitin was applied by submerging 6- to 8-week-old leaves treatment in AXS solution supplemented with or chitin for 1–2 h (control leaves submerged in AXS solution that did not contain chitin) or by feeding 6- to 8-week-old leaves with AXS solution supplemented with chitin through their petioles for 2–4 h (control leaves fed AXS without any chitin). The composition of the AXS solution composition and the petiole-feeding treatment are described in the Measurement of Kleaf and Gas Exchange section below. Prior to the submerged-leaf treatment, the leaves were incubated overnight in AXS solution to give the wound signal time to subside.

Real-time PCR was performed using Thermo Scientific ABsolute Blue qPCR SYBR Green ROX Mix in a Rotor-Gene Q (QIAGEN) machine. The program used was as follows: pre-incubation at 95 C for 15 min, followed by 40 cycles of denaturation at 95 C for 15 s each, annealing at 59 C for 30 s and extension at 72 C for 30 s. All samples were analyzed in three to four technical repetitions and at least three biological repetitions. *UBQ5* (TAIR: AT3G62250) was used as the reference gene. Relative gene expression was analyzed using the 2^-ΔΔCt^ method ^57^ The primers used are listed in **Supplementary Table 2**.

### Stomatal Aperture Measurements

For the measurements of stomatal aperture, peeled epidermal strips were prepared from the abaxial side of fully expanded leaves from 6- to 8-week-old Arabidopsis plants. The strips were immediately transferred (keeping adaxial surface facing up) to a six-well cell culture dish filled with stomatal-opening induction solution (20 mM KCl, 1 mM CaCl2, 5 mM MES; pH 6.15 adjusted with KOH) ^58,59^ and incubated under bright light (150□μmol□m^-2^□s^-1^) for 1.5□h. Then, the samples were treated with chitin (with the control left untreated) and incubated under light for an additional 1.5□h. After that treatment, the epidermal peels were placed on glass cover slips and photographed with 400X total magnification under a bright-field inverted microscope (1M7100; Zeiss) equipped with a HV-D30 CCD camera (Hitachi). Stomatal images were analyzed to calculate the aperture size using the ImageJ software (http://rsb.info.nih.gov/ij/) fit-line tool. A microscopic ruler (Olympus) was used for size calibration.

### Measurement of *K*_leaf_ and Gas Exchange

For the measurement of *K*_leaf_ and gas exchange, fully expanded leaves of similar size (approx. 6 cm ^2^) with similar vascular areas and no noticeable injuries or anomalies were harvested from 6- to 8-week-old Arabidopsis plants in the dark and immediately immersed (petiole-dipped) in artificial xylem sap (AXS; 3 mM KNO_3_, 1 mM Ca(NO_3_)_2_, 1 mM MgSO_4_, 3 mM CaCl_2_, 0.25 mM NaH_2_PO_4_, 90 μM EDFC and a micro-nutrient mix of 0.0025 μM CuSO4 * 5 H_2_O, 0.0025 μM H2MoO4, 0.01 μM MnSO4, 0.25 μM KCl, 0.125 μM H_3_BO_3_*3 H_2_O, 0.01 μM ZnSO_4_ * 7 H_2_O) with or without chitin in 0.6-mL centrifuge tubes. The tubes containing the leaves were placed in 3.3-L (20 × 20 × 8.25 cm) transparent, covered boxes containing moist tissue paper to maintain high humidity. Each box held six to eight leaf samples and served as one block in the randomized block design. The leaf samples were incubated for 2–4 h in the closed boxes under continuous light before measurements were taken.

Before any measurements were taken, each box was opened for 3 min, in order to normalize the relative humidity in the box with that of the ambient atmosphere. *K*_leaf_ was measured using the detached-leaf approach ^37^. Briefly, each leaf went through two consecutive measurements: transpiration and Ψ_leaf_. Transpiration, E, was measured using a Li-Cor 6800 gas-exchange system (Li-Cor) equipped with a 6-cm ^2^-aperture standard leaf cuvette. The measuring conditions in the cuvette were 150 μmol m^-2^ s^-1^ light with 400 μmol mol^-1^ CO_2_ surrounding the leaf, a leaf temperature of approximately 22°C and a vapor pressure deficit of approximately 1.4 kPa. The measuring conditions were set similar to the growth room conditions. Ψ_leaf_ was measured using an Arimad 3000 pressure chamber (MRC) and a homemade silicon adaptor specially designed for Arabidopsis petioles, in order to fit the O-ring of the pressure chamber. After placing the leaf in the chamber and tightly closing it, N2 pressure was gradually applied until a droplet of xylem sap was observed at the cut with the help of a Motic SMZ-171 binocular microscope (Motic) under the illumination of a KL 200 LED light (Olympus) directed toward the petiole. K_leaf_ was than calculated for each individual leaf by dividing *E* by *Ψ*_leaf_. (In our calculations, Ψ_leaf_ = ΔΨ_leaf_, as the leaf petiole was dipped in AXS at a water potential of approximately 0.) All measurements were conducted between 10 AM and 1 PM.

### Protoplast Isolation

Protoplasts were isolated from 6- to 8-week-old plants using the rapid method ^60^. Briefly, the lower leaf epidermis was peeled off from the middle part of the leaf and the peeled leaves were cut into small squares and incubated in an enzyme solution containing 3.3% (w/w) of an enzyme mix comprised of the following enzymes in the given proportions: 0.55 g of cellulase (Worthington), 0.1 g of pectolyase (Karlan), 0.33 g of polyvinyl pyrrolidone K30 (Sigma-Aldrich) and 0.33 g of bovine serum albumin (Sigma-Aldrich). That mix also contained additional substances at the following concentrations: 10 mM KCl, 1 mM CaCl_2_, 540 mM D-sorbitol, and 8 mM MES. The mix had a pH of 5.7. After 20 min of incubation at 25°C, the leaf tissue was transferred to the same solution without the enzymes and gently shaken for 5 min or until all of the protoplasts had been released into the solution. The remaining tissue debris was removed and the remaining solution containing the protoplasts was collected into a 1.5-mL tube using a cut tip. This protoplast isolation procedure yielded a large number of protoplasts (20 million protoplasts per gram of leaf tissue) in about 45 min.

### Protoplast Transformation

Protoplast complementation studies were conducted on protoplasts isolated from the leaves of *lyk5* plants. Two plasmids were co-expressed in the protoplast, one plasmid with the *AtLYK5* gene (using pML-BART-35S::AtLYK5 constructs) and a second plasmid with the GFP gene (pSAT1-EGFP-C1) ^54^ to act as a marker for a successful transformed protoplast. We co-transformed the protoplasts using polyethyleneglycol (PEG) 4000 (Sigma-Aldrich) as described by ^61^. Briefly, protoplasts were washed twice with W5 solution (155 mM NaCl, 125 mM CaCl_2_, 5 mM KCl, 2 mM MES; pH 5.7) and then resuspended in MMg solution (0.4 M sorbitol, 15 mM MgCl_2_, 4 mM MES; pH 5.7) and incubated on ice for 30 min. After that incubation, the plasmids and PEG solution (4 g PEG4000, 3 mL ddH_2_O, 2.5 ml 0.8 M sorbitol, 1 ml 1 M Ca(NO_3_)_2_) were added and incubated for 25 min at room temperature. Protoplasts were washed with W5 solution twice and then re-suspended gently in 0.5 mL of W5 and incubated in a 24-well plate at room temperature for 16 h in the dark.

### Measurement of Osmotic Water Permeability (*P*_f_)

The isolated protoplasts from SCR::GFP and *lyk5* plants and the transiently expressed *LYK5+GFP/lyk5* protoplasts were treated with chitin (control left untreated) in the 1.5 mL centrifuge tubes for 2–4 h prior to *P*_f_ measurement. The protoplast population was screened for GFP-labeled protoplasts and unlabeled cells, as described by Shatil-Cohen et al. (2011). The measurements were done using single protoplasts of both mesophyll cells and BS cells, based on the initial (recorded) rate of their increase in volume in response to hypo-osmotic challenge (transfer from a 600-mosmol isotonic bath solution to a 500-mosmol hypotonic solution). *P*_f_ was determined using a numerical approach, specifically, an offline curve-fitting procedure using the PfFIT program, as described in detail previously ^62–64^ For a detailed video description of the measurement of Pf, please refer to Shatil-Cohen et al. (2014).

### Statistical Analysis

Statistical analysis was performed using JMP 10.0 software. Means were compared using Student’s t-test, Tukey’s HSD test, as detailed for each experiment. For Tukey’s HSD test, means were considered significantly different at *P* < 0.05.

## ACKNOWLEDGEMENTS

This research was supported by the Israel Ministry of Agriculture and Rural Development (Eugene Kandel Knowledge Centers) as part of the Root of the Matter – The Root-Zone Knowledge Center for Leveraging Modern Agriculture and by the Israel Science Foundation (ISF Grant No. 878/16). AD gratefully acknowledges the Planning and Budgeting Committee (PBC) of the Council for Higher Education (CHE), Israel, for his postdoctoral fellowship. We thank Dr. Ofir Bahar for his scientific advice and critical reading of this paper.

